# Partial restoration of mutational robustness after addition of genetic polymorphism and in the presence of sexual selection

**DOI:** 10.1101/197194

**Authors:** Caroline M. Nieberding, Gilles San Martin, Suzanne Saenko, Cerisse E. Allen, Paul M. Brakefield, Bertanne Visser

**Author notes:** **Author contributions:** CMN and PMB designed the research, CMN collected and analysed data and prepared the manuscript; SS collected data; GSM analysed the data; CEA and PMB discussed and edited the manuscript; BV prepared the manuscript. **Data availability:** Data will be made available as supplementary information.

## Abstract

The interaction between mutational (i.e. genetic) robustness, cryptic genetic variation and epistasis is currently under much debate, as is the question whether mutational robustness evolved under direct selection or as a by-product of environmental robustness. Here we report that mutational robustness was restored in a mutant line of the butterfly *Bicyclus anynana* after the spontaneous mutation, *comet*, appeared in a genetically polymorphic wild type population. The *comet* mutation modified two phenotypic traits known to be under sexual selection in this butterfly: the dorsal forewing eyespot, which is normally round, but became ‘comet’-shaped, and the androconia, the structures producing the male sex pheromone, which were reduced in size. The *comet* mutant line remained phenotypically stable for ∼7 seven years, but when outcrossed to the genetically polymorphic wild type population, the outcrossed *comet* line surprisingly recovered the wild type phenotype within 8 generations. This suggests that mutational robustness against the *comet* mutation was recovered in the *comet* outcrossed line by epistatic interactions with the genetic polymorphism originating from wild types. The extent of wild type phenotype recovery in the *comet* outcrossed line was trait- and developmental temperature-dependent, such that mutational robustness was partially recovered at high, but not at low developmental temperatures. We hypothesized that sexual selection through mate choice, which is sex-reversed between developmental temperatures in this butterfly, could produce mutational robustness at a high (but not at a low) temperature. Females are the choosy sex and exert stabilizing or directional selection on male secondary sexual wing traits but only at higher temperatures. Male mating success experiments under semi-natural conditions then revealed that males with the typical *comet* mutant phenotype suffered from lower mating success compared to wild type males, while mating success of *comet* males resembling wild types was partially restored. Altogether, we document the roles of cryptic genetic variation and epistasis in restoration of mutational robustness against a spontaneous mutation with known fitness effects, and we provide experimental evidence, for the first time to our knowledge, that sexual selection can produce mutational robustness.

## Introduction

Phenotypic variation is the raw material for selection that is ubiquitous for most traits in natural populations. The amount of phenotypic variation can, however, differ dramatically within and among populations, i.e. some traits are invariant within species while being highly variable among closely related species (Flatt 2005). There is ample evidence that the amount of phenotypic variation often does not reach its full potential (i.e. there is less variation than could be present), because phenotypes are robust to mutations or to environmental perturbations (Masel and Siegal 2009; Masel and Trotter 2010). That is, most species maintain abundant genetic variation and experience a wide range of environmental conditions, but phenotypic variation remains relatively low (Waddington 1942; Félix and Barkoulas, 2015). While environmental robustness is virtually a given, as no living system can persist without regulating its internal composition and thus needs robustness to some changes in the internal or external environment, robustness to mutations is under much debate (Siegal and Leu, 2014). One critical point to be addressed by research on phenotypic robustness is to specify the causal link(s) between mutational robustness and the presence of cryptic genetic variation and epistatic interactions on phenotype stability and evolvability (Siegal and Leu, 2014), i.e. does genetic polymorphism increase or decrease phenotype stability and its evolvability? It is, furthermore, essential to determine whether or not robustness to mutations has evolved under direct selection (Meiklejohn and Hartl, 2002; Wagner et al. 1997; Masel & Siegal 2009; Lauring et al. 2013, Siegal and Leu, 2014). Many have argued that mutational robustness can result from non-adaptive processes, such as the developmental architecture underlying traits of interest (Flatt 2005; Siegal and Leu, 2014). Others have, however, suggested that robustness to mutations can evolve in populations with large population sizes or experiencing high mutation rates in response to stabilizing selection (Wagner et al. 1997; Wilke et al. 2001; Siegal and Leu 2014). Theoretical work has indeed shown that stabilizing selection reduces phenotypic variation from one generation to the next (Lande 1980; Layzer 1980; Rice 1998; Kawecki 2000). One theoretical study showed that when sexual selection operates in populations, both stabilizing and directional selection resulting from female mate choice, can favor the evolution of mutational robustness (Fierst 2013). Experimental evidence that selection drives the evolution and maintenance of mutational robustness is, however, limited, with exception of work done on RNA viruses (Montville et al. 2005; Sanjuan et al 2007; McBride et al. 2008).

The butterfly *Bicyclus anynana* is an important model in evolutionary ecology for studies on sexual selection and developmental plasticity, including seasonal polyphenism (Brakefield et al. 2009). Several wing traits were shown to play an important role in sexual selection, including the UV-reflecting white pupils of dorsal forewing eyespots (Costanzo and Monteiro 2007; Prudic et al. 2011) and the male sex pheromone produced partly by male-specific wing structures called androconia (Costanzo and Monteiro, 2006; Nieberding et al., 2008; San Martin et al., 2011). Behavioral experiments manipulating these traits in males showed that females exert stabilizing sexual selection on males for round-shaped and small to mid-sized pupils (Robertson and Monteiro 2005), as well as directional sexual selection on increasing quantities of male sex pheromone components (Nieberding et al. 2012; van Bergen et al. 2013). Frankino et al (2005) also revealed stabilizing selection on wing size, although that may be due to natural selection on flight ability and not female choice. Developmental temperature generates morphologically distinct seasonal forms adapted to either the wet or the dry African seasons (Brakefield et al. 2009). This is adaptive phenotypic plasticity as the wet and dry phenotypes are produced non-randomly with respect to high or low developmental temperatures. Sexual roles are reversed across wet and dry seasons: while females exert choosiness and males compete for accessing mates during the wet season, males become the choosy sex during the dry season (Prudic et al. 2011). *Comet* is a spontaneous, recessive and pleiotropic mutation that arose in a single individual of the *B. anynana* wild type population before 1998 (Brakefield 1998; Brakefield and French 1999; Beldade et al. 2009). The *comet* (*cc*) mutation produces several large phenotypic changes on the wing traits that affect male mating success. Namely, the dorsal forewing eyespot is pear-shaped (“comet-shaped”) instead of round, and the androconia are either reduced in size on the forewing or absent on the hindwing (Brakefield, 1998; Brakefield and French, 1999; Fig 1). The *comet* mutant line displayed a stable phenotype in the laboratory for at least seven years, while reared at various developmental temperatures (Brakefield et al. 1998; Brakefield and French 1999; Brakefield 2001).

**Fig 1:**
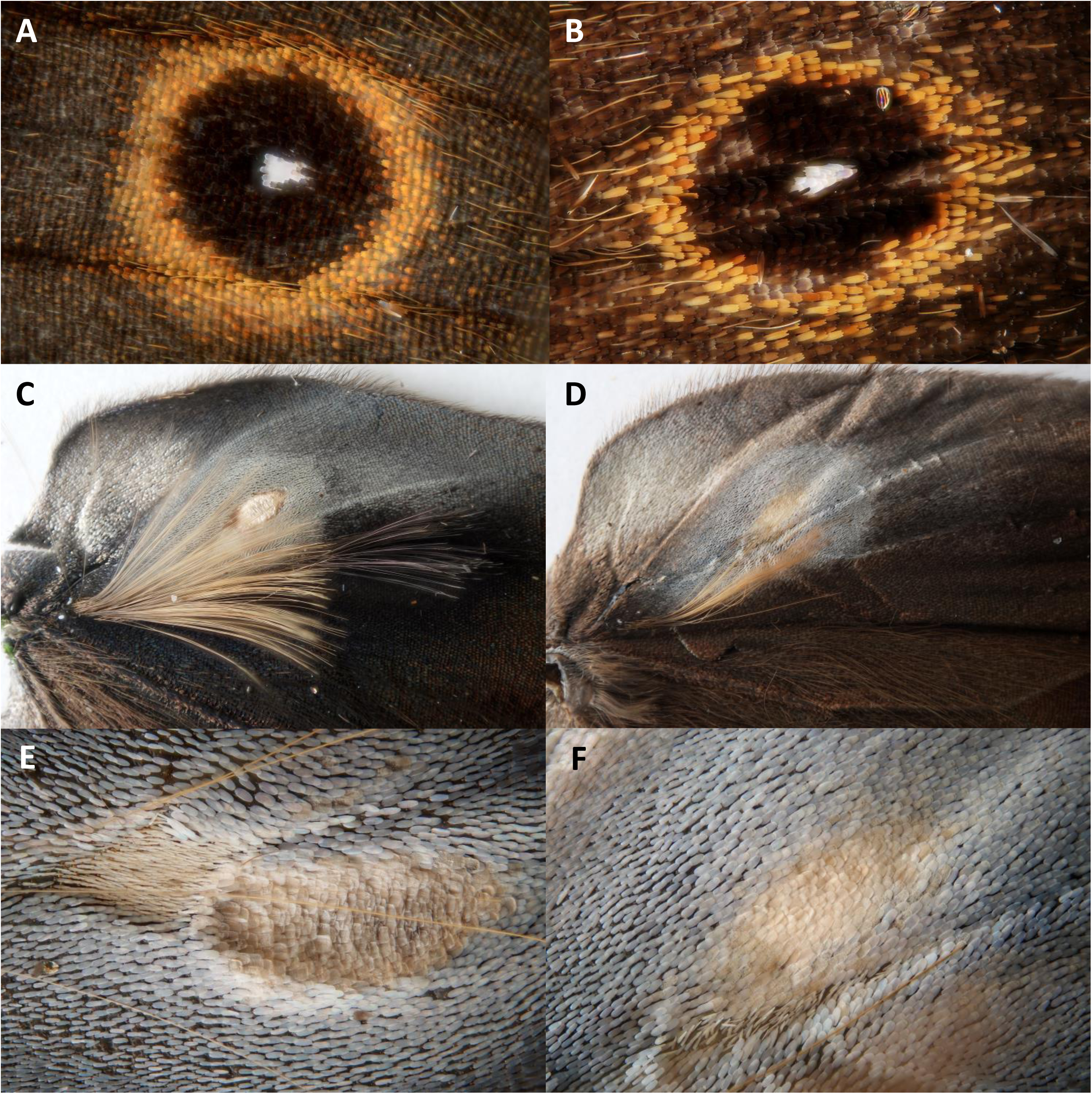
Morphological differences between wild type (left) and *comet* (right) individuals: posterior eyespot on the dorsal side of the forewing (A, B), androconial first and second hairpencils along with the second androconial patch on the dorsal side of the hindwing (C, D), and detail of the second androconial patch (after removal of the hair pencils; E, F).

In this study, we outcrossed the *comet* inbred line to the wild type population that displays high levels of heterozygosity (Van’t Hof et al. 2005) in order to restore the genetic polymorphism typical of the wild type population around the *comet* mutation. Surprisingly, in the next few generations we observed that most individuals of the outcrossed *comet* line that were reared at 27°C degrees and developed in the wet seasonal form, did not express the *comet* phenotype and could not be distinguished from wild types. Loss of the *comet* phenotype in the outcrossed *comet* line suggested that mutational robustness against the *comet* spontaneous mutation was restored by epistatic interactions with the genetic polymorphism that was cryptic in the wild type population. Yet, when reared at 20°C and developing in the dry seasonal form, the outcrossed *comet* line again fully expressed the *comet* phenotype. Hence, mutational robustness against the *comet* mutation was dependent on the developmental environment of the outcrossed *comet* individuals. In order to document these qualitative observations, we quantified the effect of the *comet* mutation in the outcrossed *comet* line on both morphological (eyespot size and shape, androconia presence and size) and physiological (amounts of male sex pheromone components) secondary sexual phenotypic traits by comparing outcrossed *comet* with wild types reared at various temperatures typical of the dry (20°C) and wet (27°C) seasonal forms. It remained unclear, however, why mutational robustness against *comet* would be restored at a high but not at a low rearing temperature. We hypothesized that sexual selection through female preference for round-shaped and small to mid-shaped eyespot pupils and/or for large male sex pheromone quantities may have fuelled the rapid recovery of phenotypic robustness in the outcrossed *comet* line. We thus expected that sexual selection should act at the high but not at the low rearing temperature given that sexual roles are plastic and females are the choosy sex only at a high temperature (Prudic et al 2011). To test this hypothesis, we performed behavioral experiments to compare the mating success of males from the outcrossed *comet* (*cc)* line displaying the *comet* (reared at 20°C) or the wild type (reared at 27°C) phenotype with wild type (*++*) males competing for wild type (*++*) wet seasonal females reared at 27°C.

## Material and methods

### Insects

An outbred wild type population of the African butterfly, *Bicyclus anynana* (Lepidoptera: Nymphalidae), was established in 1988 from over 80 gravid females collected from a single source population in Malawi, Africa. *B. anynana* larvae were maintained on a maize-based diet (*Zea mays*), whereas adults were fed mashed banana (*Musa acuminata*). High levels of heterozygosity were maintained by using laboratory population sizes that ranged between 400 and 600 adults per generation (Brakefield 2001; Van’t Hof et al. 2005). The wild type population was reared in climate rooms at a set of different temperature (20-27°C) and humidity regimes (60 to 80% RH) that represent the natural range of environmental variation present in the field. The two extreme temperatures, 20°C (± 1°C) and 27°C (± 1°C), represent the developmental temperature typical of the dry and wet seasonal forms under laboratory conditions, respectively.

The *comet* line founded before 1998 is formed by homozygous “*cc”* individuals displaying pear-shaped (“comet-shaped”) instead of round eyespots on the dorsal and ventral sides of fore- and hind-wings (Fig. 1; Brakefield et al. 1998; Brakefield and French 1999; Beldade et al, 2009). Genetic diversity within the *comet* line is expected to be low, first due to the initial bottleneck as this spontaneous recessive mutation occurs very rarely in the wild type population, and second because the *comet* line was subsequently kept in the laboratory at a relatively small population size for years. In this study, we thus restored the genetic diversity at loci other than *comet* was restored. The collected F_1_ generation (*c+*) displayed a wild type phenotype and was crossed among itself to produce a F_2_ generation in which ¼ of the individuals displayed the *comet* phenotype and were “*cc*”, similarly to findings in Beldade et al (2009). These F_2_ *comet* “*cc”* individuals were selected to produce the next generations of what we call hereafter the “outcrossed *comet* line”.

### Effect of *comet* mutation on male wing secondary sexual traits

To quantify the phenotypic effect of the *comet* mutation and assess the effect of developmental temperature on its expression, we reared 3 wild type and 8 *comet* families obtained from eggs collected in the outcrossed *comet* line about 6 to 8 generations after the F_2_ generation at 5 temperatures: 19, 21.5, 23, 24.5 and 27°C. Eggs were collected from the outcrossed *cc* line and from the wild type population. We measured the following male traits: (i) pupil length/width ratio of the dorsal forewing posterior eyespot pupil (measured as the maximal length of the pupil parallel to the wing vein and the width as the maximum width perpendicular to the length), (ii) pupil area of the dorsal forewing posterior eyespot pupil (approximated from the area of an ellipse with pupil length as major axis and pupil width as minor axis), (iii) the area of the first androconial patch located on the forewing ventral side, (iv) the area of the second androconial patch located on the hindwing dorsal side, (v) the presence/absence of a well-developed hairpencil (functionally associated with the forewing androconia), and (vi) presence/absence of a well-developed hairpencil (associated with the hindwing androconia). Hairpencils were considered to be well-developed when at least 10 hairs were present. These six morphological traits are either directly or indirectly (i.e. androconia size) involved in sexual selection (Nieberding et al. 2012; Bacquet et al. 2015). We also estimated the area of the forewing and hindwing by measuring the area between 4 landmarks on each wing. For all morphometric measurements, we recorded the x y coordinates of different landmarks by projecting an image of each morphological structure of interest from a stereomicroscope equipped with a camera lucida onto a graphical tablet. The x y coordinates were then converted into areas or lengths taking into account the magnification and the number of pixels between the coordinates.

### Effect of *comet* mutation on male sex pheromone quantities

Eggs were collected from the outcrossed *comet* line (*cc*) 6 to 8 generations after the F_2_, and from the wild type population. Individuals were kept at 20°C or 27°C throughout development and adult life. Virgin males were sampled for determining male sex pheromone (MSP) quantities at ages 3, 7, 14 and 21 days for individuals kept at 27°C and ages 3, 7, 14 and 28 days for individuals kept at 20°C. MSPs were extracted and quantified as described previously (Nieberding et al. 2008). Briefly, one forewing and hindwing per individual were soaked during 5 minutes in 600µl of hexane, after which 1 ng/µl of internal standard (palmitic acid) was added. Extracts were then analyzed on a Hewlett-Packard 6890 series II gas chromatograph (GC) equipped with flame-ionization detector and interfaced with a HP-6890 series integrator with nitrogen as carrier gas. The injector temperature was set at 240°C and the detector temperature at 250°C. A HP-1 column was used and temperature increased from the initial temperature of 50°C by 15°C/min up to a final temperature of 295°C, which was maintained for 6 min.

### Effect of *comet* phenotype on male mating success

To test for behavioral effects of the *comet* mutation on male mating success, we performed behavioural experiments competing wild type (++), heterozygote (*c*+) and outcrossed *comet* (*cc*) males for mating success. Two behavioral experiments were performed that aimed at comparing mating success of wild type males and *comet* males that showed both abnormal pupil shapes and lacked androconia (experiment 1), or *comet* males that had normal pupil shapes but lacked androconia (experiment 2). Specifically, for experiment 1: wild type males were obtained from eggs of the wild type stock population; *comet* males were obtained from the outcrossed *comet* line (F_3_ generation); heterozygote males (c+) by crossing 32 F_2_ *cc* virgin females from the outcrossed *comet* line with 30 wild type males, and 30 F_2_ *cc* males from the outcrossed *comet* line with 28 virgin wild type females, in two separate cages. Eggs of the three treatments (cc, c+ and ++) were collected for 10 days and reared mostly at 27°C, although eggs from replicates 2 and 3 of experiment 1 were kept at the beginning of their development at 20°C in order to delay emergence of the adults.

We noted that about 10% of *cc* males (60 out of 600 males) in the outcrossed *comet* F_3_ generation displayed wild type eyespots, while androconia remained typically “*comet*-like” with the second set of hairpencils being reduced. To test how reduced androconia alone (with normal eyespots) affected male mating success, we crossed these 60 *comet* F_3_ males with 50 *comet* F__3__ females that had also more rounded eyespots to produce the F_4_ generation of the *comet* outcrossed line, which were used in experiment 2. The F_4_ generation of the outcrossed *comet* line produced mostly males with a wild type eyespot shape but *comet*-like reduced androconia. We compared the mating success of these F_4_ outcrossed *comet* males with that of male heterozygote (c+) and wild type (++) males obtained as described above for experiment 1.

In both behavioral experiments, groups of 3 to 10-day old virgin males were released in a spacious tropical greenhouse that provided a semi-natural environment for *B. anynana*. Male genitalia were dusted with colored fluorescent powder (Joron and Brakefield 2003; Nieberding et al. 2008). In experiment 1, males (*cc, c+* and *++*) competed for matings at a 1:1:1 ratio, with group numbers ranging from 60 to 75 males per group. In experiment 2, wild type (*++*), heterozygote (*c+*) and *comet* (*cc*) males were released in a proportion of 1:1:2 to mimic an environment in which the wild type phenotype (represented by both *++* and *c+* males) was as abundant as the *comet* phenotype, with numbers ranging between 25 to 60 males per group. In both experiments, 3 to 10-day old virgin wild type females (50 to 130 per replicate) were released the following morning, to obtain approximately a 2:1 male:female ratio. Males competed for matings during 72 h, after which females were inspected under ultraviolet illumination for fluorescent dust transferred during mating to assess female mate choice. Double matings occurred occasionally (approximately 1 in every 20 matings) and were scored as 1:1. Experiment 1 was repeated three times, and experiment 2 was repeated twice.

### Statistics

All statistical analyses were performed with R 2.12.0 (R Development Core Team 2010), using the lme4 package (Bates et al. 2015). To test for effects of *comet* on wing morphology, we used mixed models with family as a random variable, and type (outcrossed *comet* or wild type), temperature (as continuous variable) and their interaction as fixed explanatory variables. We used a normal error distribution for the continuous variables androconial patch area and eyespot pupil size, and a binomial distribution for the hairpencils, which were scored as present or absent. Eyespot pupil ratio data was log transformed and pupil surface was square root transformed to improve homoscedasticity and normality of residuals. For model parameter inference, we used Markov Chain Monte-Carlo simulations (i.e. the mcmc function from the lme4 package) for normal models and approximate z tests for binomial models. Temperature values were centered on the maximum value (27°C). The "type effect" parameter, therefore, corresponds to the difference between *comet* and wild type at 27°C. For pupal and androconial patch size, wing area (centered on the mean) was also added as an explanatory variable to control for wing size.

To analyze sex pheromone quantities, individuals reared at 20°C and 27°C were analyzed separately, because selected age classes differed between the two temperatures. We used linear models with MSP titers as dependent variables and age, type (outcrossed *comet* or wild type) and their interaction as explanatory variables. These explanatory variables were tested with type II F tests (nested models comparison, with main effects tested after removing their interaction from the full model).

To analyze effects of the *comet* mutation on male mating success, replicated G tests of goodness of fit were used as described by Sokal & Rohlf (1995). A single G test of goodness of fit was computed for each replicate independently and three additional G statistics were calculated: a heterogeneity G test to test whether the different replicates show the same trend, a pooled G test based on the pooled dataset for all replicates and a total G test based on the sum of the single G statistics produced for each replicate.

## Results

### Effect of *comet* mutation on male wing secondary sexual traits

Within 6-8 generations following outcrossing, we quantified the phenotypic traits affected by the *comet* mutation by comparing eyespot shape and size as well as androconia size between families from the wild type population and from the outcrossed *comet* line. When reared at 27°C, the phenotypes of the outcrossed *comet* line had almost completely recovered the wild type phenotype: eyespot pupil shape (circular compared to the elongated pupils of the original inbred *comet* line), size of the second androconial spot, and presence of the first androconial hairpencil were similar between outcrossed *comet* families and wild type families (Fig 2, Table 1). In contrast, phenotypes of outcrossed *cc* families displayed increasing differences compared to the wild type when developmental temperature was decreased (Fig 2, Table 1). Thus, several generations after outcrossing, the effect of the *comet* mutation had become strongly temperature-dependent for all traits except the size of the second androconial patch, and the effect of the mutation was uncoupled across the set of six affected traits (Table 1).

**Table 1:** Model estimates for 6 male morphological traits involved in sexual selection from wild type and *comet* mutants (type) across 5 breeding temperatures. The four first models are linear mixed models with family as random effect (not shown) and normal error distribution. The inference on model parameters is based on 10000 MCMC simulations. The two last models (presence/absence of well-developed hairpencils) are generalized linear mixed models with family as random effect (not shown), binomial error distribution and logit link function. The inference on parameters is based on approximate z tests. The temperature values are centered on 27°C so that type effect estimates the difference between *comet* and wild type at 27°C. (^+^) The type x temperature interaction was not significant for the first androconial hairpencil (p>0.99), but this model had some estimation problems due to the high proportion of "presence" in wild type individuals; yet the graphs (Fig 2 panels E, F) show that the models provide a good fit of the data and that there is no doubt that the differences observed between *comet* and wild type for the androconial hairpencils depend on temperature (i.e. significant interaction) too.

| Posterior pupil surface (mm<sup>2</sup>, square root transformed) | Estimate | Std.Error | 95% credible interval | p |  |
| --- | --- | --- | --- | --- | --- |
| Intercept | 0.7756 | 0.0304 | [0.7124 ; 0.8374] | 0.0000 | *** |
| forewing surface (mm<sup>2</sup>, centered on the mean) | 0.0063 | 0.0012 | [0.0040 ; 0.0087] | 0.0000 | *** |
| type (stock) | -0.2017 | 0.0671 | [-0.3379 ; -0.0676] | 0.0058 | ** |
| temperature (°C, centered on 27°C) | -0.0154 | 0.0037 | [-0.0228 ; -0.0080] | 0.0000 | *** |
| type * temperature | 0.0446 | 0.0105 | [0.0239 ; 0.0656] | 0.0000 | *** |
| Posterior pupil length/width (log transformed) | Estimate | Std.Error | 95% credible interval | p |  |
| Intercept | -0.0084 | 0.0318 | [-0.0726 ; 0.0558] | 0.7806 |  |
| type (stock) | 0.0811 | 0.0777 | [-0.0750 ; 0.2391] | 0.3030 |  |
| temperature (°C, centered on 27°C) | -0.0780 | 0.0046 | [-0.0871 ; -0.0689] | 0.0000 | *** |
| type * temperature | 0.0603 | 0.0138 | [0.0327 ; 0.0876] | 0.0000 | *** |
| First androconial patch surface (mm<sup>2</sup>) | Estimate | Std.Error | 95% credible interval | p |  |
| Intercept | 0.5576 | 0.0206 | [0.5161 ; 0.5994] | 0.0000 | *** |
| hindwing surface (mm<sup>2</sup>, centered on the mean) | 0.0032 | 0.0007 | [0.0018 ; 0.0045] | 0.0000 | *** |
| type (stock) | 0.1037 | 0.0422 | [0.0234 ; 0.1867] | 0.0160 | * |
| temperature (°C, centered on 27°C) | 0.0160 | 0.0022 | [0.0117 ; 0.0204] | 0.0000 | *** |
| type * temperature | -0.0308 | 0.0057 | [-0.0421 ; -0.0198] | 0.0000 | *** |
| Second androconial patch surface (mm<sup>2</sup>) | Estimate | Std.Error | 95% credible interval | p |  |
| Intercept | 1.0250 | 0.0341 | [0.9579 ; 1.0915] | 0.0000 | *** |
| forewing surface (mm<sup>2</sup>, centered on the mean) | 0.0094 | 0.0011 | [0.0073 ; 0.0115] | 0.0000 | *** |
| type (stock) | 0.0350 | 0.0692 | [-0.1020 ; 0.1677] | 0.5984 |  |
| temperature (°C, centered on 27°C) | 0.0580 | 0.0033 | [0.0514 ; 0.0647] | 0.0000 | *** |
| type * temperature | -0.0111 | 0.0088 | [-0.0289 ; 0.0057] | 0.2050 |  |
| Presence of well-developed first hairpencil | Estimate | Std. Error | z value | p |  |
| Intercept | 4.6489 | 0.6301 | 7.3778 | 0.0000 | *** |
| type (stock) | -1.8899 | 1.2852 | -1.4706 | 0.1414 |  |
| temperature (°C, centered on 27°C) | 0.7554 | 0.0968 | 7.8064 | 0.0000 | *** |
| type * temperature | -4.8537 | 432.6316 | -0.0112 | 0.9910<sup>+</sup> |  |
| Presence of well-developed second hairpencil | Estimate | Std. Error | z value | P |  |
| Intercept | -0.9744 | 0.3671 | -2.6545 | 0.0079 | ** |
| type (stock) | 4.5342 | 1.4172 | 3.1994 | 0.0014 | ** |
| temperature (°C, centered on 27°C) | 0.6429 | 0.1514 | 4.2467 | 0.0000 | *** |
| type * temperature | -0.54 | 0.2825 | -1.9113 | 0.0560 | (*) |

**Fig 2:**
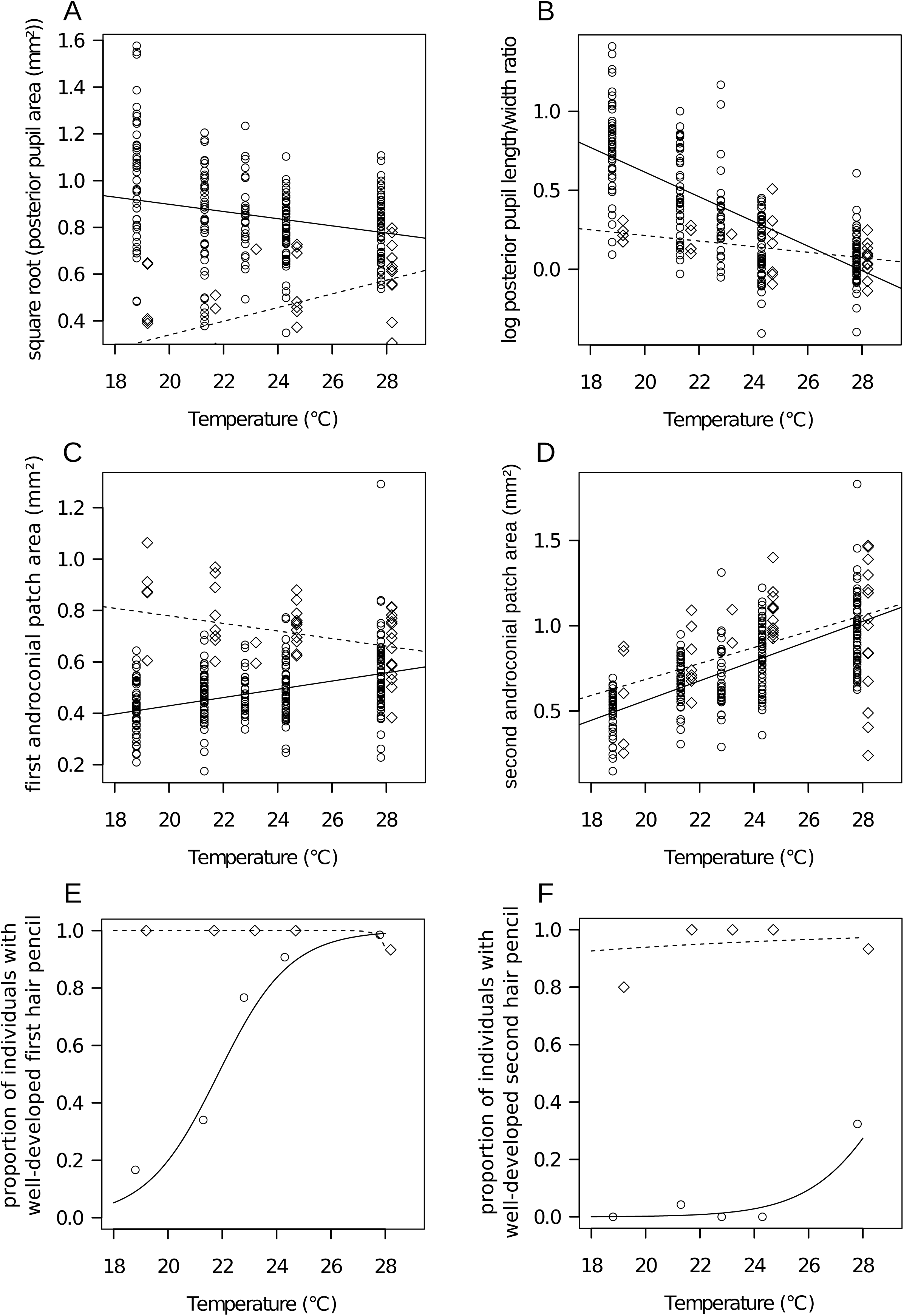
Six male morphological traits were measured for wild type stock (diamonds) and *comet* mutants (circles) across 5 breeding temperatures, including (A) the posterior pupil area in mm^2^ (square root transformed), (B) the posterior pupil length/width ratio (log transformed), (C) the first androconial spot area (mm^2^), (D) the second androconial spot area (mm^2^), (E) the proportion of individuals with a well-developed first hairpencil, and (F) the proportion of individuals with a well-developed second hairpencil. The lines represent the corresponding mixed model predictions for wild type (dotted lines) and *comet* (continuous lines) which were corrected for wing size on graphs A, C and D.

### Effect of *comet* mutation on male sex pheromone quantities

The quantities of male sex pheromone (MSP) components were compared between wild type and *comet* males randomly chosen from the outcrossed *comet* line to test if morphological changes induced by the *comet* mutation affected MSP production. MSP production did not differ between wild type and outcrossed *comet* males reared at a higher temperature, but differed strongly at the lower temperature. At 27°C, titers of MSP1, MSP2 and MSP3 of wild type and outcrossed *comet* males displayed a similar pattern across age classes (Fig 3; none of the age x type interactions were significant at the 0.05 level: MSP1: F=1.1, df=3, p=0.35 - MSP2: F=1.02, df=3, p=0.38 - MSP3: F=1.83, df=3, p=0.14). In stark contrast, patterns of MSP titers differed strongly between *comet* and wild type males when butterflies were reared at 20°C. The production of MSP2 was almost completely suppressed in all ages in outcrossed *comet* males reared at 20°C, due to absence of the androconia (Fig 3; age x type interaction: F=11.02, df=3, p<0.0001). Additionally, MSP1 and MSP3 titers of outcrossed *comet* and wild type males both progressively increased, but peaked at 28 days of age in outcrossed *comet* males versus 14 days of age in wild type males. MSP1 and MSP3 titers subsequently decreased in wild type males (age x type interaction MSP1: F=10.79, df=3, p=0.001; MSP3: F= 10.75, df=3, p=0.005) (Fig. 3). MSP1 and MSP3 titers at a single age class (14-day old) in outcrossed *comet* males were similar to MSP titers of the younger age class in wild type males (8-day old). Thus the rate of increase of MSP1 and MSP3 titers was slower in outcrossed *comet* than in wild type males at 20°C.

**Fig 3:**
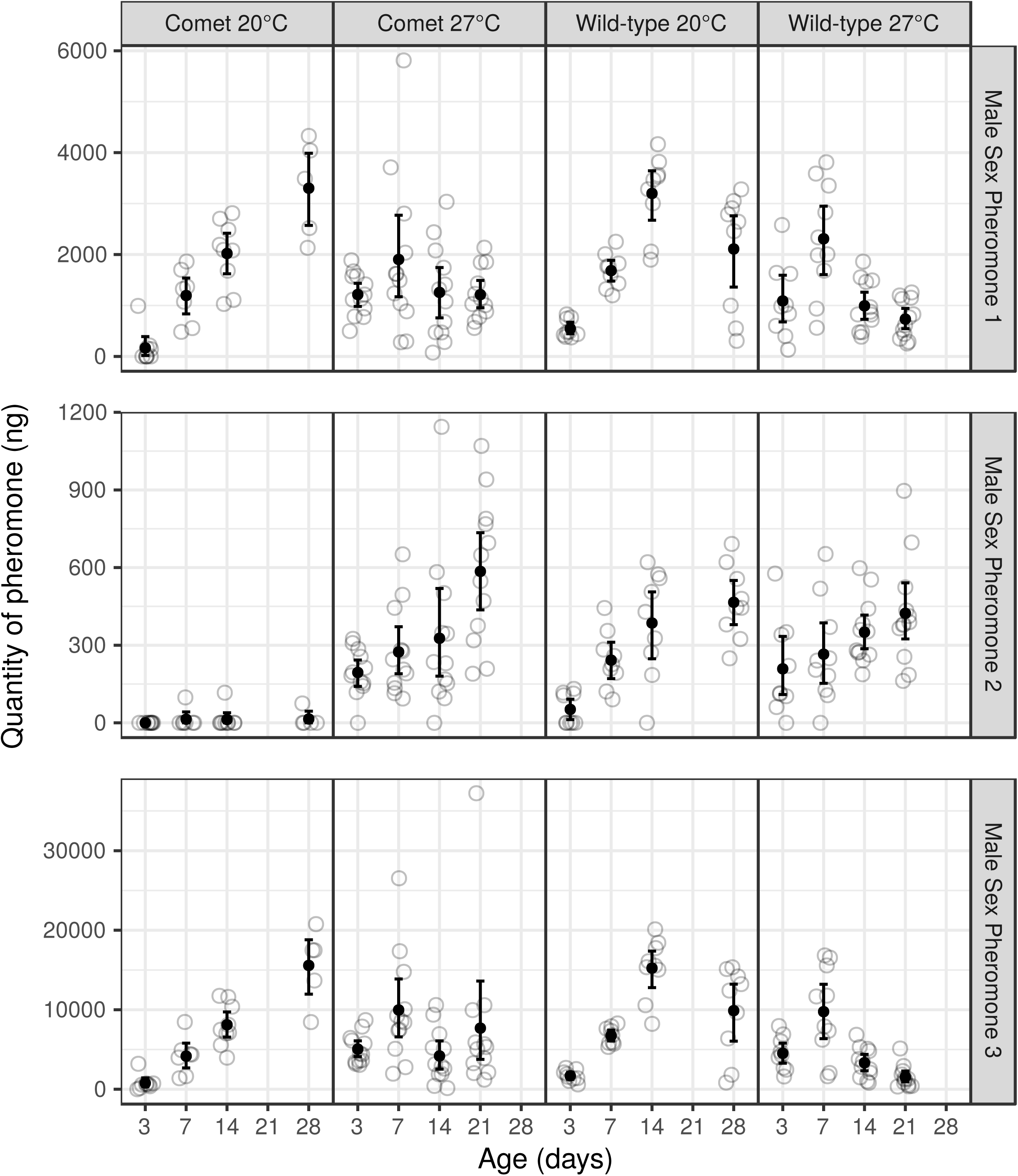
Male sex pheromone (MSP; top = MSP1; center = MSP2; bottom = MSP3) titers of *comet* (cc) and wild type (++) males reared at 20°C and 27°C and sampled at 5 different ages (3, 7, 14, 21, 28). At 20°C, no individuals were sampled at 21 days, while at 27°C, and no individuals were sampled at 28 days. The open gray circles show the observed values, the black dots represent their mean and the error bars represent bootstrap 95% confidence intervals for the mean. Some random noise has been added on the x axis to limit overplotting.

### Effect of comet phenotype on male mating success

Mating success of males with *comet* or wild type phenotypes was compared during two mating competition experiments under semi-natural conditions in a large tropical greenhouse. In the first experiment, we used outcrossed *comet* males of the F_3_ generation, most of which (540/600) had reduced androconia and modified eyespot pupils typical for *comet* mutants (Brakefield, 1998; Brakefield and French 1999). Mating success of outcrossed *comet* males (*cc*) was significantly lower than that of heterozygote (*c+*) or wild type males (*++*) and was similar for all three replicates: both the total and pooled G-tests were significant, as well as the single G-tests for two out of three replicates (Table 2). During the second experiment only outcrossed *comet* males with circular-shaped eyespot pupils and reduced androconial hairpencils from the F_4_ generation were selected to compete for matings. Mating success of outcrossed *comet* (*cc*) males was significantly lower than that of heterozygote (*c+*) or wild type (*++*) males in the first replicate, but not in the second replicate, with non-significant pooled and global G-tests (Table 2). In both experiments, outcrossed *comet* (*cc*), heterozygote (*c+*) and wild type (*++*) males were recaptured in similar proportions to those at which they were released (all G-tests were non-significant at the 0.05 level; Table 2); hence male survival was similar among competing groups of males.

**Table 2:** A. Male recapture rates in behavioral experiments and replicated G tests for goodness of fit for each experiment. B. Male mating success in behavioral experiments and replicated G tests for goodness of fit for each experiment.

| **A. Male recapture rate - First experiment** |  |  |  |  |  |  |
| --- | --- | --- | --- | --- | --- | --- |
| ++ | *c+* | *cc* | *type of G test* | *df* | *G* | *P* |
| 33 | 18 | 22 |  | 2 | 4.82 | 0.0899 |
| 14 | 19 | 21 |  | 2 | 1.49 | 0.4742 |
| 42 | 45 | 33 |  | 2 | 2.00 | 0.3675 |
|  |  |  | Total | 6 | 8.31 | 0.2160 |
| 89 | 82 | 76 | Pooled | 2 | 1.03 | 0.5983 |
|  |  |  | Heterogeneity | 4 | 7.29 | 0.1215 |

| **Male recapture rate - Second experiment** |  |  |  |  |  |  |
| --- | --- | --- | --- | --- | --- | --- |
| ++ | *c+* | *cc* | *type of G test* | *df* | *G* | *p* |
| 18 | 21 | 32 |  | 2 | 0.92 | 0.6306 |
| 16 | 16 | 34 |  | 2 | 0.06 | 0.9701 |
|  |  |  | Total | 4 | 0.98 | 0.9124 |
| 34 | 37 | 66 | Pooled | 2 | 0.31 | 0.8567 |
|  |  |  | Heterogeneity | 2 | 0.67 | 0.7141 |

| **B: Male mating success - First experiment** |  |  |  |  |  |  |  |
| --- | --- | --- | --- | --- | --- | --- | --- |
| ++ | *c+* | *cc* | *type of G test* | *df* | *G* | *p* | *sig.* |
| 23 | 18 | 5 |  | 2 | 13.22 | 0.0013 | ** |
| 12 | 15 | 4 |  | 2 | 7.18 | 0.0276 | * |
| 14 | 12 | 6 |  | 2 | 3.54 | 0.1706 |  |
|  |  |  | Total | 6 | 23.93 | 0.0005 | *** |
| 49 | 45 | 15 | Pooled | 2 | 22.02 | 0.0000 | *** |
|  |  |  | Heterogeneity | 4 | 1.91 | 0.7527 |  |

| **Male mating success - Second experiment** |  |  |  |  |  |  |  |
| --- | --- | --- | --- | --- | --- | --- | --- |
| ++ | *c+* | *cc* | *type of G test* | *df* | *G* | *p* | *sig.* |
| 13 | 11 | 9 |  | 2 | 7.24 | 0.0268 | * |
| 5 | 6 | 11 |  | 2 | 0.09 | 0.9555 |  |
|  |  |  | Total | 4 | 7.33 | 0.1193 |  |
| 18 | 17 | 20 | Pooled | 2 | 4.17 | 0.1242 |  |
|  |  |  | Heterogeneity | 2 | 3.16 | 0.2059 |  |

## Discussion

This study provides an example of the restoration of mutational robustness against a spontaneous mutation that has detrimental effects on secondary sexual phenotypic traits. *Comet* mutants have appeared sporadically in the wild type laboratory-reared population of *Bicyclus anynana* (Brakefield 1998; Brakefield, 2001; Beldade et al. 2009). Individuals with this recessive, pleiotropic mutation deviate from wild types in a number of wing characteristics, including eyespot and androconial sizes and shapes, which are of critical importance for male mating success (Robertson and Monteiro 2005; Costanzo and Monteiro 2006; Nieberding et al. 2008, 2012). Mutant phenotypes were stable at a range of developmental temperatures for at least 7 years following isolation of the mutant from the wild type population (Brakefield 1998, 2001; Brakefield and French 1999). While a quarter of the F_2_ generation issued from the crossing between heterozygote (*c+*) individuals displayed, as expected, the *comet* phenotype, the latter faded away within the next few generations within the outcrossed *comet* line when individuals were reared at a high developmental temperature. Our question was why phenotypes returned to wild type values, and why this happened only at the higher developmental rearing temperature.

Over the years significant progress has been made in understanding the molecular basis of phenotypic robustness to genetic perturbations. Epistatic interactions, molecular chaperones (such as HSP90), and functional redundancy by gene duplications play a primary role in maintaining genetic robustness (Rutherford and Lindquist 1998; Hartman et al. 2001; Landry et al. 2007; Levy and Siegal 2008; Siegal and Leu 2014; Fares 2015). In our documented case with *comet*, genome size and structure are conserved in the *comet* mutant line before and after the cross with the wild type population; hence gene duplications nor epistatic interactions alone were responsible for restoration of the wild type phenotype in the *comet* outcrossed line.

### Role of genetic polymorphism in mutational robustness

We suggest that the addition of genetic polymorphism to the genetic background of the *comet* mutation through outcrossing with wild types allowed the partial restoration of mutational robustness against *comet*. Genetic polymorphism and epistatic interactions between the *comet* allele and wild type alleles at other loci can thus have provided the raw material for selection to favor mutational (i.e. genetic) robustness. While the evolutionary relevance of cryptic genetic variation for adaptation is generally accepted (e.g. Hayden et al. 2011), the role of cryptic genetic variation in producing or maintaining mutational robustness remains unclear (Siegal and Leu, 2014). A few pieces of experimental evidence have recently emerged. First, robustness to mutations in the P450 protein was higher in larger and more polymorphic populations compared to smaller and less polymorphic populations such that genetic polymorphism is responsible for higher mutational robustness (Bloom et al. 2007). It was also shown that the genetic background in which the HSP90 chaperone is expressed can have a large effect on resultant phenotypes, as is the case in *Drosophila* (Rutherford and Lindquist 1998). How would genetic polymorphism restore mutational robustness of the *comet* mutant line? The genetic polymorphism that was added in the outcrossed *comet* line is cryptic in the sense that wild type individuals did not show phenotypic variation for the traits affected by the *comet* mutation. Most mutations are background-dependent (i.e. show epistatic effects) and this cryptic genetic variation accumulated in the wild type population such that it caused diversification of genetic backgrounds with which the *comet* mutation interacted epistatically. Some genetic backgrounds of the wild type population produced a particular phenotypic effect with the *comet* mutation and others not. The diverse genetic background of the wild type population would then lead the *comet* outcrossed line as a whole to express more new phenotypes than when it was inbred (Wagner 2007, 2011, 2012; Siegal and Leu 2014). This conceptual argument positively correlates mutational robustness with evolvability, which has been formalized in mathematical models of so-called neutral networks in genotype space (more recently termed genotype networks) and has some empirical support (McBride et al. 2008, Hayden et al. 2011, Lauring et al. 2013, in Siegal and Leu 2014). Our results thus suggest that genetic polymorphism might be required for phenotypic robustness to be restored although large and robust empirical evidence is, to the best of our knowledge, currently missing.

### Role of sexual selection in mutational robustness

Importantly, in our *comet* case study the genetic polymorphism present in the wild type population was not sufficient to *maintain* mutational robustness - otherwise the *comet* mutant would not have appeared in the wild type population in the first place. Genetic polymorphism merely allowed partial *restoration* of mutational robustness after polymorphism was added to the original *comet* mutant line. This suggests that genetic polymorphism alone was not sufficient to maintain mutational robustness. As the *comet* mutation affects secondary sexual traits which we know are under sexual selection in *B.anynana*, we suggest that sexual selection against the *comet* phenotypic led to the restoration of mutational robustness against the mutation. In the wild type *B. anynana* population the eyespot and androconial traits that are affected by the *comet* mutation are known targets of sexual selection. Here, we showed that wild type males had higher mating success compared to *comet* males, and moreover, that outcrossed *comet* males with ‘less extreme’ (i.e., closer to the wild type) phenotypes have higher mating success than males with more extreme phenotypes. For outcrossed *comet* males with less pronounced eyespot and androconial deformations, mating success was higher than the outcrossed males with more pronounced changes in eyespot and androconial traits. Decreased mating success of these outcrossed *comet* males may be due to their larger (compared to wild type) eyespot pupils (Robertson and Monteiro 2005), and reduced MSP transfer to female antenna during courtship as a consequence of the reduced second hairpencil and androconial spots (Nieberding et al. 2008). MSP amounts presents on male wings are indeed correlated with androconia spot areas (Nieberding et al. 2012). Strong sexual selection on phenotypically diverse outcrossed *comet* males likely led to very rapid allelic changes at loci other than *comet*, which interacted epistatically with the *comet* mutation to produce more wild type phenotypes.

The importance of sexual selection in driving trait evolution has long been recognized (Andersson et al. 1998), but it has remained elusive whether selection, including sexual selection, could play a role in evolving phenotypic robustness. Sexual section is often assumed to be a directional force triggering the evolution of exaggerated traits (i.e. traits with disproportionate scaling), but sexual selection can also be a stabilizing force that either rapidly increases, or reduces, differentiation in male traits over generations. A first modeling study by Fierst (2013) suggested that female mate preferences increase male phenotypic robustness under three different sexual selection scenarios compared to a randomly mating population. Her theoretical results imply that female choice leads to selection pressures that affect mutational robustness, which thus has the potential to develop in any population experiencing sexual selection (Fierst 2013). Our results suggest that sexual selection restored mutational robustness against the spontaneous *comet* mutation within a few generations at high rearing temperature, likely by stabilizing selection for *comet* phenotypic variants that were closer and closer to the wild type trait values. To the best of our knowledge we provide the first experimental evidence suggesting sexual selection may act as a driver for restoring mutational robustness.

The *comet* phenotype was originally temperature-independent and pleiotropically affected several sexually selected traits. Outcrossed *comet* individuals displayed phenotypic plasticity for trait expression in response to temperature and the expression of several abnormal traits became uncoupled. At 27°C outcrossed *comet* males recovered the first androconial hairpencil and formed an eyespot similar in shape to that of wild type males. This observation of trait- and temperature-specific recovery of mutational robustness in outcrossed *comet* mutants excludes the possibility that the *comet* mutation was lost as a consequence of introducing the wild type background. A study on *D. melanogaster* revealed that insertional mutations of 16 genes led to temperature-dependent phenotypic effects on wing size, where no differences were found at 18°C, but smaller wing sizes were found at 27°C (Debat et al. 2009). Moreover, both mutations and temperature affected the level of fluctuating asymmetry, where fluctuating asymmetry in shape remained unaffected by temperature, but individual variation became apparent at 19°C. Mutational robustness in these mutants may thus be less efficient at a lower temperature, similar to our findings in *B. anynana*. The partial recovery of phenotypic robustness and preservation of the *comet* phenotype at the lower temperature can be explained by two non-exclusive hypotheses. First, mutational robustness can be limited in more stressful environments (de Visser et al. 2003; Fares 2015) and a lower temperature is indeed more stressful for this tropical butterfly (Brakefield et al. 2007; Steigenga and Fischer 2009). Second, differences in sexual selective pressures at 20 and 27°C may account for the preservation of the *comet* phenotype at the lower temperature. A study by Prudic et al (2011) showed that sexual selection was plastic in *B. anynana*: while males are under strong competition for mating by choosy females at 27°C, it is the males that become the choosy sex at 20°C. Females may, therefore, have induced directional selection for *comet* males displaying normally shaped eyespot pupils and increased MSP2 titers (correlated to larger androconial structures), but only at 27°C. Based on the results of our behavioural experiments, we suggest that female preference for wild type eyespot and hairpencil characters was the basis for strong sexual selection on *comet* modifier loci that were introduced into the population through outcrossing with wild type individuals, and that this selection brought the *comet* mutation under the control of a temperature-dependent genetic switch. This switch thus suppresses many aspects of the *comet* phenotype at 27°C, but not at 20°C.

It is important to note one weakness of our work, which is that we tested sexual preferences of wild type and not of *comet* females, where we assumed that both would have similar preferences for male traits. This may not be true, because, for example, learning through imprinting of male phenotypes during sexual maturation is known to affect female sexual preferences in insects, also in *B. anynana* (e.g. Westerman et al, 2012). *Comet* females may thus have learned to prefer the male *comet* phenotype because they grew up together. Learning is biased in *B. anynana*, however, in that females can learn to prefer supra-natural sexual stimuli, but not reduced wing ornamentation and thus females may not be able to learn to prefer drab *comet* males (Westerman et al, 2012). Assortative mating with similar phenotypes may also affect sexual preferences of *comet* females toward *comet* male phenotypes although we have no evidence for assortative mating in *B. anynana*. It is important to note, however, that we observed no restoration of phenotypes for the *comet* mutation during the 7 years the inbred *comet* line was kept in the laboratory when *comet* females had no choice to mate with other males than phenotypic *comet* ones.

In conclusion, this study provides, to the best of our knowledge, a first empirical example that suggests that genetic polymorphism and sexual selection can underlie the rapid evolution of increased phenotypic robustness of abnormal phenotypes towards wild types. We documented a fortuitous example where cryptic genetic variation had been decoupled from the arising of a new spontaneous mutation: genetic polymorphism present in the wild type was added to the mutant isogenic line after this mutation was observed to be stably present. This approach, following the fate of spontaneous mutations decreasing mutational robustness after adding different levels of genetic polymorphism around the mutation, could be useful to implement as a novel method to experimentally assess the effect of background genetic polymorphism on the restoration of mutational robustness in more natural settings, as evolution proceeds.

## Acknowledgments

This work was supported by the Fonds de la Recherche Scientifique - FNRS under grant no; 24905063 and 29109376. This is BRC publication 408 of the Biodiversity Research Centre.

## Notes

**Conflict of interest:** The authors declare no conflict of interest.

